# On the Boundary Conditions of Avoidance Memory Reconsolidation: An Attractor Network Perspective

**DOI:** 10.1101/826909

**Authors:** Rodrigo MM Santiago, Adriano BL Tort

## Abstract

The reconsolidation and extinction of aversive memories and their boundary conditions have been extensively studied. Knowing their network mechanisms may lead to the development of better strategies for the treatment of fear and anxiety-related disorders. In 2011, Osan et al. developed a computational model for exploring such phenomena based on attractor dynamics, Hebbian plasticity and synaptic degradation induced by prediction error. This model was able to explain, in a single formalism, experimental findings regarding the freezing behavior of rodents submitted to contextual fear conditioning. In 2017, through the study of inhibitory avoidance in rats, Radiske et al. showed that the previous knowledge of a context as non-aversive is a boundary condition for the reconsolidation of the shock memory subsequently experienced in that context. In the present work, by adapting the model of Osan et al. (2011) to simulate the experimental protocols of Radiske et al. (2017), we show that such boundary condition is compatible with the dynamics of an attractor network that supports synaptic labilization common to reconsolidation and extinction. Additionally, by varying parameters such as the levels of protein synthesis and degradation, we predict behavioral outcomes, and thus boundary conditions that can be tested experimentally.

## Introduction

Much of the brain functioning can be understood from the connectionist perspective, where mental phenomena are described by the synaptic interconnection between neural networks (Feldman and Ballard, 1982). Studies suggest that memories are encoded from the synaptic configuration of neuronal assemblies (Choi et al., 2018), where patterns of experience-dependent activity are stored and retrieved by the association of sensory information (Stein, 1998). This is possible due to the plastic aspect of the synapses, which can constantly be strengthened or weakened by biochemical processes depending on whether the activity of the involved neurons correlates or not (Hebb, 1949; Hughes, 1958; Neves et al., 2008). Neural networks thus evolve by associating the activation of certain engrams to recurrent patterns of sensory signals and internal processes (Gonzalez et al., 2019). In turn, synaptic conformations form flexible dynamic systems that tend to guide the flow of information, characterizing the so-called attractor networks (Amit, 1992; Rolls, 2007; Wills et al., 2005).

In 2011, Osan et al. modified the Hopfield neural network model of associative memory (Hopfield, 1982) by adding the possibility of synaptic degradation by prediction error (Pedreira et al., 2004; Sevenster et al., 2013). In addition, the model allows the modulation of Hebbian learning (Hebb, 1949), simulating the blockade of protein synthesis through amnesic drugs, and the modulation of the synaptic degradation itself, simulating the effect of drugs acting on its blockade or reinforcement (Alberini, 2009; Kandel, 2001; Lamprecht and LeDoux, 2004). This new model maintains the characteristics of a connectionist model and a nonlinear dynamic system where the neural states vary in the time domain toward points of convergence called attractors, i.e., local minima in an energy base that represents the network state-space (Mizusaki et al., 2016; Seung, 1996; Zylberberg and Strowbridge, 2017). The retrieval of a memory pattern (attractor) previously stored can be achieved from the partial or noisy knowledge of its content (Ritter et al., 1992).

Empirical evidence demonstrates that the dynamics of the population activity of neurons from regions such as the hippocampus and neocortex behaves as attractor networks (Brunel, 2016; Pfeiffer and Foster, 2015; Rolls, 2007). For example, the associative characteristic of the hippocampus, mainly because of the highly connected structure of recurrent collaterals in CA3 (Amaral and Witter, 1989; Ishizuka et al., 1990; Rennó-Costa et al., 2014), allows it to rapidly encode activity from the cortex, and consequently retrieve them from sensory cues. Thus, declarative and context-fearful memories, dependent on the medial temporal lobe, are coded faster than the others (Henke, 2010; Marr et al., 1991; Treves and Rolls, 1994). Before coming to CA3, the afferent signal from the entorhinal cortex goes through a pattern separation process in the dentate gyrus (Drew et al., 2013; Sahay et al., 2011), allowing different representations between similar experiences and, thus, minimizing interference in the storage and retrieval of specific memories (Yassa and Stark, 2011).

Memories stabilized via the consolidation process (Kitamura et al., 2017; McGaugh, 2015) can be updated from their reactivation, when they become momentarily labile by the induction of new intracellular biochemical cascades (Bevilaqua et al., 2008; Lee, 2009; Nader et al., 2000; Tronson and Taylor, 2007). Such a phenomenon, known as reconsolidation, restabilizes memory traces, leading to structural modifications and synaptic configuration updating (Lee et al., 2006, 2017; Tronson et al., 2006). Depending on the animal’s environmental perception leading to memory retrieval, the related engram can be reinforced from, for example, gene expression and protein synthesis *de novo* (Milekic and Alberini, 2002; Nader et al., 2000; Suzuki et al., 2004), or degraded via the ubiquitin-proteasome pathway (Ehlers, 2003; Hegde et al., 1993; Kaang et al., 2009; Lee et al., 2008) and autophagy (Shehata et al., 2018). Moreover, the association of the synaptic degradation with the formation of a new memory trace related to the perception of a safe environment may result in the extinction phenomenon, when the animal extinguishes the original behavioral reaction against a particular cue or context and starts to exhibit a new behavioral response related to the activation of the novel engram (Izquierdo et al., 2004; Pavlov, 1927; Rescorla and Heth, 1975). In other words, the degree of synaptic degradation, and consequently the occurrence of reconsolidation or extinction, is associated with the prediction error level, which measures the degree of mismatch between the retrieved memory and the engram activated by the current event (Sevenster et al., 2014).

The circumstances that allow the occurrence of memory reconsolidation or extinction are known as boundary conditions, defined as physiological, environmental or psychological variables that may influence the behavioral phenotype (Fernández et al., 2016; Nader and Hardt, 2009), delimiting situations in which the memory becomes unstable, i.e., subjected to changes. Therefore, investigating the boundary conditions leading to the reconsolidation or extinction of aversive memories is especially important to help developing better strategies for the treatment of anxiety and fear-related disorders (Debiec and Ledoux, 2004; Kindt et al., 2009; Lee et al., 2005). Some already known boundary conditions are the strength of the aversive memory storage (Forcato et al., 2014; Suzuki et al., 2004; Taylor et al., 2009; Wang et al., 2009), the time elapsed between the storage and the aversive memory reactivation (Baratti et al., 2008; Forcato et al., 2013; Inda et al., 2011; Milekic and Alberini, 2002), and, more recently, the previous knowledge of the context/stimulus as non-aversive (Radiske et al., 2017). However, the results of the literature that define them are controversial (Nader and Hardt, 2009), suggesting the necessity of a broad parametric mapping that adequately defines the actual existence of the boundary conditions already discovered.

In order to map the structural biological parameters with the boundary conditions of memory reconsolidation, we investigated the network mechanisms involved in the reconsolidation of aversive memory in the inhibitory avoidance (IA) paradigm, also known as avoidance memory (Ader et al., 1972; Boccia et al., 2004; Wilensky et al., 2000). To that end, we considered the computational approach of the attractor network model developed by Osan et al. (2011), which was previously used to explain experimental findings regarding freezing behavior of rodents submitted to paradigms of contextual fear memory consolidation, reconsolidation, and extinction. Here, we use such a model to gain theoretical insights into the experimental results of Radiske et al. (2017), which showed that the prior non-aversive learning of the context where the animals experience the shock (i.e., the IA box) is a boundary condition for the reconsolidation of the avoidance memory.

## Methods

### Code development

The attractor network and all simulations were implemented using Python 2.7 and the SciPy scientific package (Jones et al., 2001). The code is available online at https://github.com/tortlab/attractor-network.

### Attractor network

The attractor network developed by Osan et al. (2011) is an adaptation of the Hopfield neural network — an associative memory system, also known as content-addressable memory (Hopfield, 1982). In the new model, neural activity varies between 0 and 1 (instead of -1 and +1 as in the original Hopfield network). This simple change avoids the storage of opposite memory patterns by eliminating connection symmetry between two neurons and the strengthening of connections between inactive units. In this way, the model allows a more realistic representation of both neuronal activity (firing rate) and synaptic connections (Osan et al., 2011).

All simulations were performed in networks composed of *n* = 100 neurons, where the activity *u* of each unit *i* was calculated by the following differential equation:

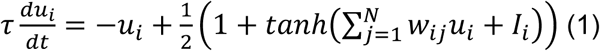

where *τ* = 1 is a constant that determines the update rate of the state *u*_*i*_, *w*_*ij*_ is the synaptic weight of the connection between the presynaptic neuron *j* and the postsynaptic neuron *i*, and *I* is the network input signal that corresponds to external and internal information that influences the animal’s perception of the environment (Osan et al., 2011). The equation was solved by Euler’s method with 100 integration points over time from *t* = 0 to *t* = 10.

Connections between units are organized in the *n*x*n*-dimensional matrix *W*, where positive and negative values represent the strength of excitatory and inhibitory synapses, respectively. Synaptic weights are updated through a process consisting of two terms (*n*x*n* matrices *HLP* and *MID*, see below) dependent on the steady-state **u** (*n*-dimensional vector) achieved after a strong input **I** (*n*-dimensional vector with values varying in [-5,+5]). Therefore, the process of memory retrieval is inserted into the storage process by directly influencing both terms that determine the variation of *W*:

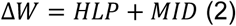

where *HLP* refers to Hebbian Learning Plasticity, and *MID*, to Mismatch-Induced Degradation (Osan et al., 2011).

The *HLP* term is calculated by *HLP* = *S*(**u**\***u**^**T**^) - *S*((1-**u**)\***u**^**T**^), where the asterisk represents the outer product and *S* ≥ 0 corresponds to the level of Hebbian plasticity that covers requirements such as receptor activation, intracellular signaling, and protein synthesis. The connection between a presynaptic neuron *j* and a postsynaptic neuron *i* is reinforced by *S* if *u*_*i*_ = *u*_*j*_ = 1, in case of maximum mutual activation. If *u*_*j*_ = 1 and *u*_*i*_ = 0, then the connection is changed by -*S*. And if *u*_*j*_ = 0, independent of *u*_*i*_, there is no impact on the connection. Intermediate effects occur in the case of intermediate values of *u*_*j*_ and *u*_*i*_.

The *MID* term is calculated by *MID* = *D*(**m**\***u**^**T**^), where the synaptic degradation factor *D* ≥ 0 represents the level of protein degradation. *MID* depends on the level of the prediction error, represented by **m** (*n*-dimensional vector), which is determined by the mismatch between the normalized input **I** (**I**_**norm**_ ∈ [0,1]) and the retrieved state **u**, that is: **m** = **I**_**norm**_ – **u**. Thus, *MID* is a nonzero matrix if **u** diverges from **I**_**norm**_, and acts towards weakening afferent connections responsible for the mismatch.

The values of the synaptic weights are truncated at a minimum of -1 and a maximum of +1. Finally, synaptic weights are subject to a time-dependent decay:

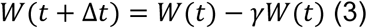

where *γ* ∈ [0,1] determines the decay rate.

Figure 1A shows the flow of the memory storage process, where the numbers inside the circles represent the equations above. For the retrieval tests, the flow process is shown in Figure 1B, where the weak input **I** corresponds to the context signal (see below) with values varying between 0 and 0.1.

**Figure 1.**
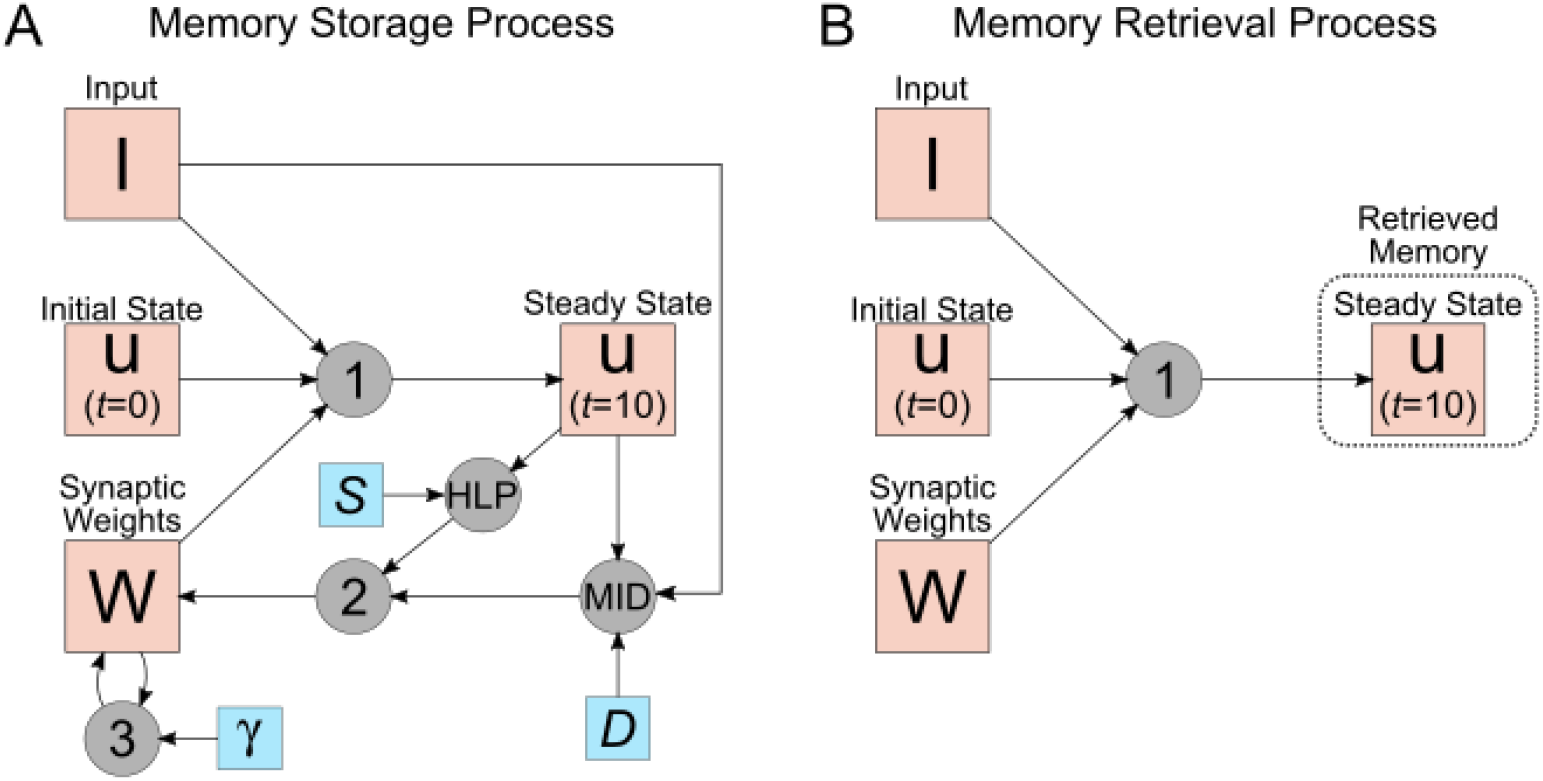
Schematics of memory storage and retrieval processes in the network. A) The synaptic weight matrix is updated depending on the steady-state activity reached by the network following a strong input signal, the initial network state, and a previous synaptic configuration. Two main factors determine the weight changes: *S* simulates the protein synthesis level and *D*, the synaptic degradation level. The former follows a Hebbian plasticity rule and the latter is dictated by the amount of mismatch between the input and the steady state. The synaptic weight matrix is also updated by the time-dependent decay factor γ. B) The retrieved memory is represented by the network steady state that is achieved upon a weak input, the initial network state, and the current synaptic configuration. The big squares represent vectors or matrices, where **I** is the external input signal applied to the network, **u** is the network state and **W** is the synaptic weight matrix. The small squares represent factors (i.e., parameters) belonging to the respective equations represented by circles. *S*: protein synthesis factor; *D*: degradation factor; γ: time-dependent decay factor. HLP: Hebbian Learning Plasticity equation; MID: Mismatch-Induced Degradation equation; Numbers correspond to the equations in Methods.

### Experimental protocol and network input patterns

To simulate the main experimental protocol of Radiske et al. (2017), we used the schema shown in Figure 2A, where each session corresponds to the respective input pattern shown in Figure 2B. The first stage is the storage of the non-related memory by the input **I**_**nr**_ in all networks (notice that the activated neurons in this pattern do not overlap with those in the other patterns). Then, in the next stage, two “habituation” groups are distinguished: one that receives the control memory (**I**_**c**_) simulating exploration of an open field, and the other, the non-shock memory (**I**_**ns**_), simulating the exploration of the IA box without shock. As in previous work (Osan et al., 2011), for the latter memory a set of neurons representing the context is activated (yellow square) along with another set of neurons representing the absence of danger. Subsequently, the networks undergo the training session, corresponding to the simulation of the IA task and the storage of the shock (“avoidance”) memory (**I**_**s**_). In this memory pattern, the same context neurons as in **I**_**ns**_ are activated along with a different set of neurons representing the presence of danger (Osan et al., 2011). The last storage occurs at the nonreinforced reexposure session with a pattern equivalent to a mixture of shock and non-shock memories (**I**_**r**_). Each habituation group (control and non-shock) is then subdivided into two others: “vehicle”, simulating the animals that received saline shortly after reexposure and have normal Hebbian learning; and “aniso”, simulating networks with inhibited protein synthesis (or a related effect) due to injection of an amnesic drug, such as anisomycin. Retrieval tests may be performed after each memory storage session (see below).

**Figure 2.**
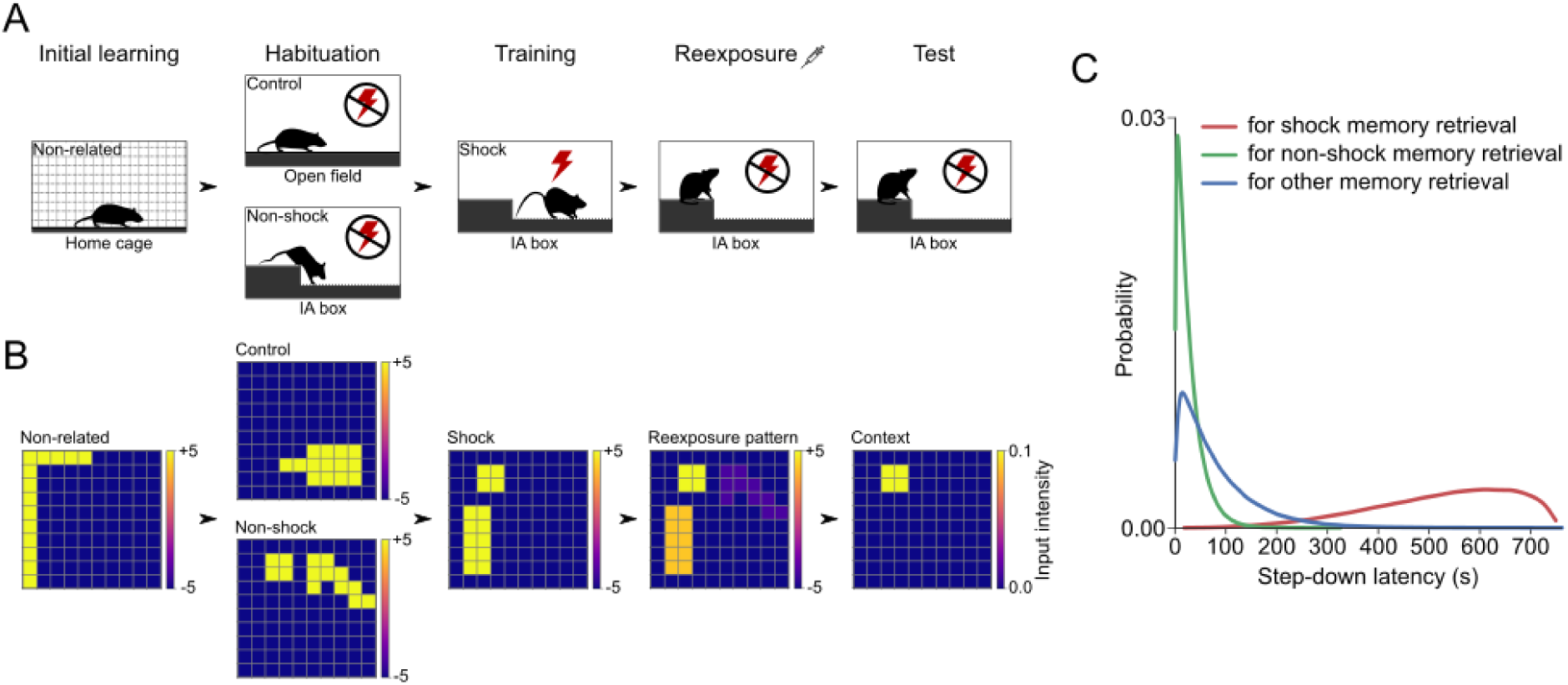
Experimental protocol, network input patterns and behavioral response probability. A) Schematics depicting the simulation stages (see text), which are related to the main experimental protocol of Radiske et al. (2017). Although only represented as the last step, retrieval tests can be performed after any storage stage. B) Input patterns (**I**) associated with each session in A. Positive and negative values represent excitatory and inhibitory input signals, respectively. The reexposure input pattern was meant to simulate a conflictual environment perception from the 40 seconds that the animals explore the platform without stepping down (Radiske et al., 2017). In this input pattern, the shock neurons are more excited than the non-shock ones (see Methods). C) Probability density functions for step-down latency in retrieval tests according to the retrieved memory pattern. Shock memory retrieval is associated with long latencies, while the retrieval of other memories leads to short latencies.

**Figure 3.**
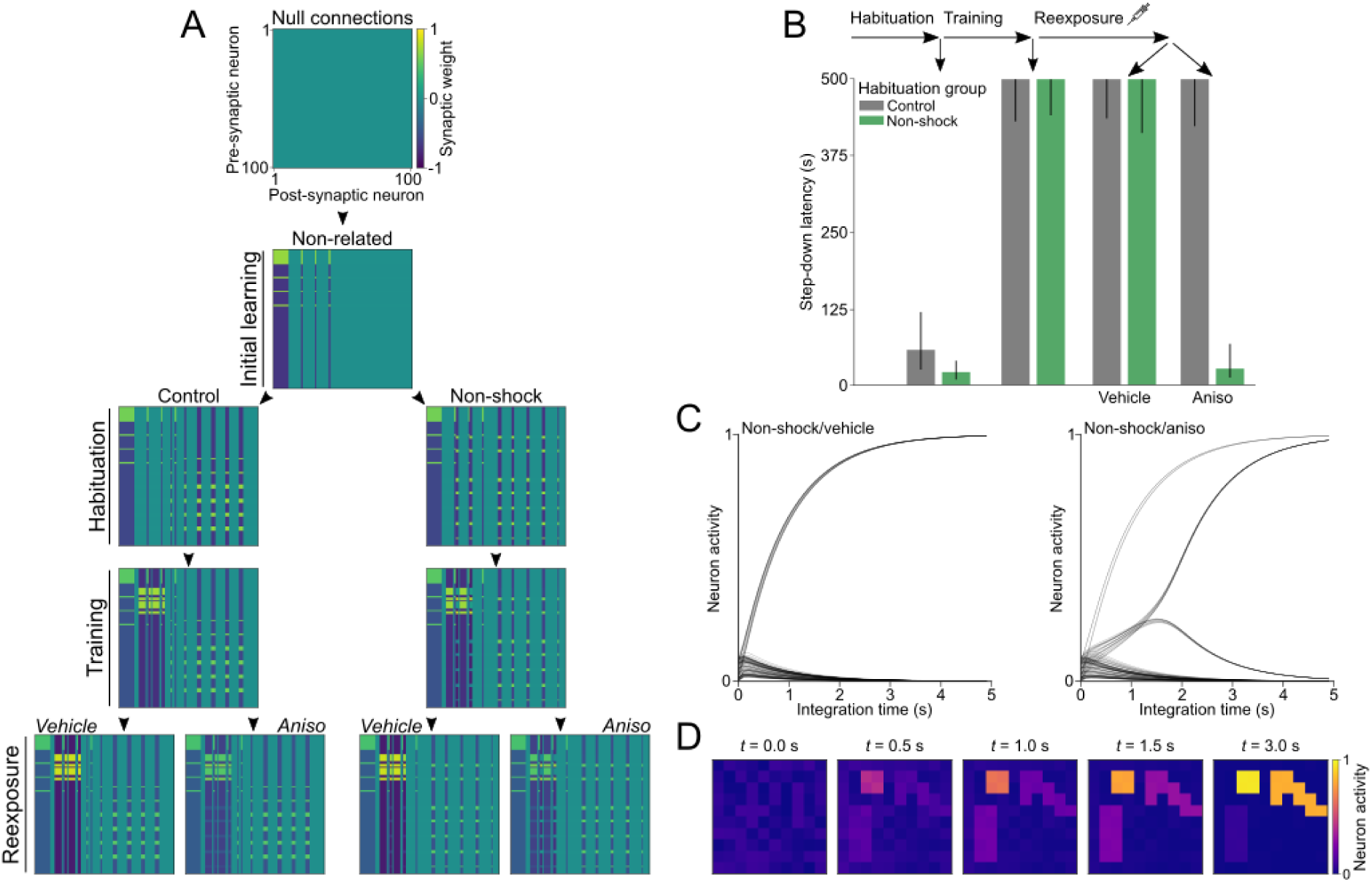
Synaptic configuration and retrieval tests throughout the experimental protocol. A) Matrices corresponding to the configuration of synaptic weights after each session of memory storage. From a null matrix, the initial learning process stores the non-related (e.g., home cage) memory pattern (*S* = 0.8). In the habituation session, two groups are distinguished by the storage of control or non-shock memories (*S* = 0.8). After that, both groups store the shock pattern in the training session (*S* = 0.85). Thereafter, vehicle and aniso groups are defined by the level of Hebbian plasticity following the reexposure session (*S* = 0.85 for vehicle and *S* = 0 for aniso). To simulate the interval of 1 day between each session, the synaptic matrix was updated using a decay rate of γ = 0.15. Positive and negative values represent excitatory and inhibitory synapses, respectively. B) After each storage stage (horizontal arrows), 1000 retrieval tests were simulated (vertical/diagonal arrows) and the results are presented as the median ± interquartile range. Notice that reconsolidation blockade only occurs for the aniso networks of the non-shock habituation group. C) Examples of non-shock/vehicle (left) and non-shock/aniso (right) network dynamics during retrieval. The network activity of the first group converges to the shock pattern, while the activity of the second network initially wavers between the shock and non-shock patterns, and then converges to the latter one. D) Frames of the non-shock/aniso network dynamics depicted in C. Despite the initial increasing activity of the shock neurons, the non-shock attractor is retrieved in the steady state.

To simulate the 24h interval between one session and another, the matrix of synaptic weights was updated after the storage of each memory via equation 3, with *γ* = 0.15. The storage strength itself depended on the session: *S* = 0.8 at initial storage and habituation; *S* = 0.85 at training; at reexposure, *S* = 0.85 for vehicle and *S* = 0 for aniso; and *D* = 1.25 at all sessions, except during parametric analysis of such factor. The higher *S* values in the training and reexposure sessions emulate the emotional nature of the aversive memory (as opposed to the neutral, non-related memory), that is, the involvement of the amygdala in strengthening the (re)consolidation of the shock experience.

For the retrieval tests, the initial network state was randomly assigned using a uniform distribution between 0 and 0.1 to set the activity of each neuron, except for the context neurons which receive a weak input fixed at 0.1. Note that context neurons are those active both in the shock and non-shock memory patterns and are meant to represent common aspects of the experience (e.g., the box appearance). For each simulation, 1000 retrieval tests were performed and their results presented as the median with interquartile range. For the color plot graphs, each point is the median of 1000 tests.

The input pattern **I**_**r**_ during reexposure corresponds to a mixture of the shock and non-shock memories, as shown in Figure 4A. Specifically, it is defined by **I**_**r**_ = **I**_**s**_ + (**I**_**ns**_ -**I**_**s**_)*f*(#), where *#* ∈ [*#*_*min*_ = 0, *#*_*max*_ = 10] – the “pattern number” – is related to the duration of the nonreinforced reexposure session, and *f*(*#*) = 1 / (1 + *e*^(*#max*/2)-#^) is a function representing the ratio between input patterns **I**_**ns**_ and **I**_**s**_ and varies monotonically between 0 and 1 (Osan et al., 2011). To simulate a 40-second reexposure session (Radiske et al., 2017), we used # = 3.1, which corresponds to **I**_**r**_ closer to the aversive memory (Figure 2B).

**Figure 4.**
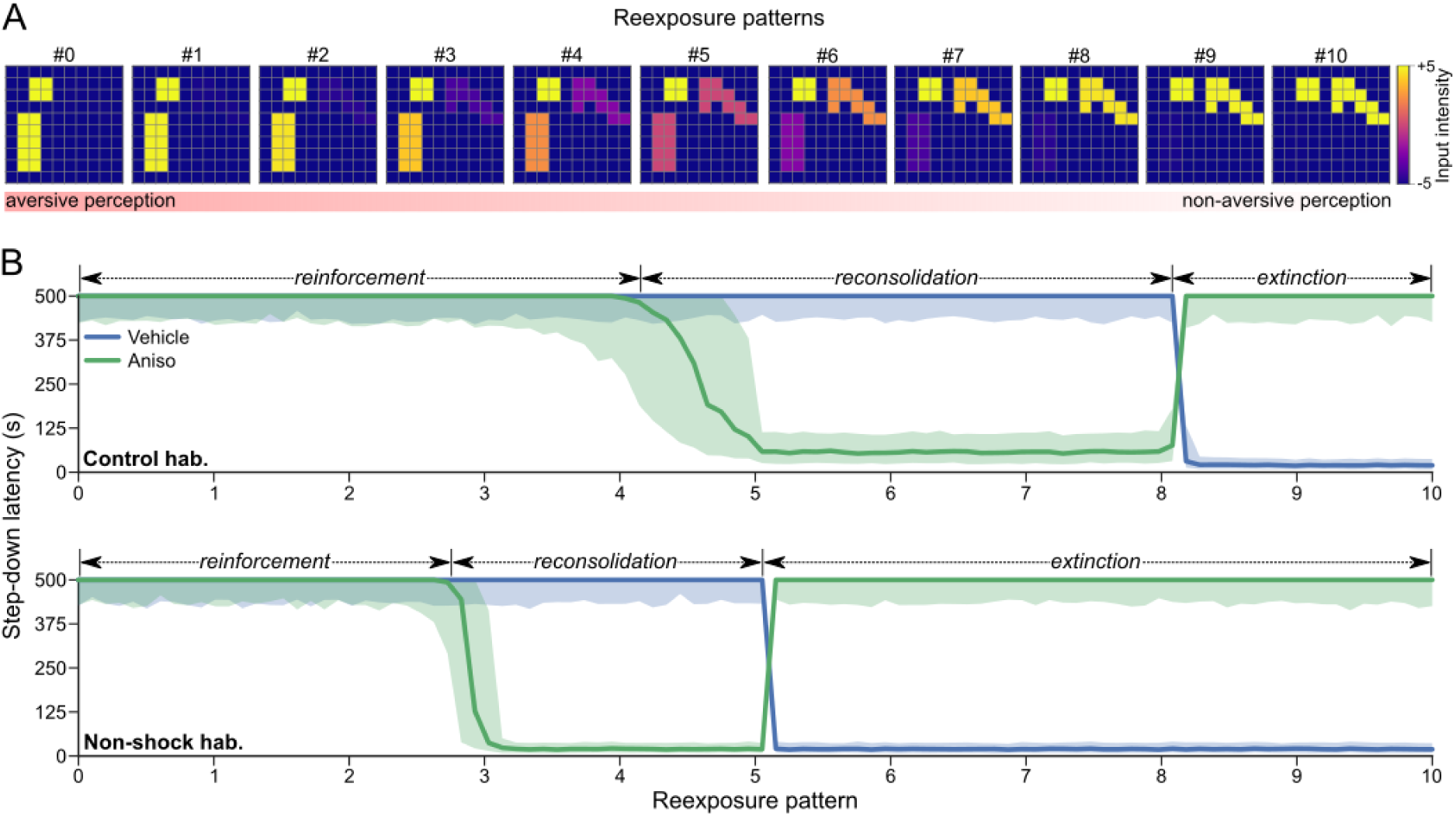
The reexposure pattern as a boundary condition for reconsolidation. A) Reexposure input pattern for different values of the parameter #, which simulates the animal’s perception of the IA box following reexposure. A change from aversive to non-aversive perception occurs with increasing this parameter (from 0 to 10), which relates to the duration of the nonreinforced reexposure. B) Step-down latencies in post-reexposure retrieval tests as a function of the reexposure input pattern #. Notice that the non-shock habituation group anticipates the reconsolidation and extinction phenomena in relation to their occurrence in the control habituation group. For example, whereas avoidance memory extinction appears from reexposure pattern #8 for the control habituation group, the same phenomenon occurs from pattern #5 for the non-shock habituation group. Results depict the median ± interquartile range of 1000 retrieval tests for each parameter choice.

### Simulation of animal behavior

In the IA task, the animal (in this case, rat) is initially placed on a platform inside a box and expected to rapidly step down to the floor level following its natural instinct to explore the whole context, unless it has previously experienced an aversive sensation in the floor, such as a foot shock. Thus, the latency for the animal to step down the platform is taken as a measure of a previously learned shock/avoidance memory. To simulate this behavior, we define three probability density functions for step-down latencies, as shown in Figure 2C. At each retrieval test, a step-down latency is chosen randomly from the probability density function corresponding to the retrieved memory pattern. If the retrieved pattern is the shock memory, the latency is more likely to be high (red curve), that is, the animal is more likely to avoid exploring the floor and stay over the platform. Otherwise, the probability is higher for low latency values. In particular, for retrieval of the non-shock memory, the probability is even higher for low latency values (green curve) since it indicates that the animal already knows the environment as non-aversive.

The probability density functions follow distributions of type *Beta*(α, β), in which *f*_shock_ = 750*Beta*(3.52,1.5) for shock memory retrieval; *f*_non-shock_ = 15000*Beta*(1.1,600) for non-shock memory retrieval; and *f*_other_ = 35000*Beta*(1.1,480) for the retrieval of the others memory patterns (Figure 2C). The latency values are truncated at the ceiling of 500 s, i.e., the maximum duration of the reexposure session in Radiske et al. (2017).

## Results

In order to investigate the boundary condition reported by Radiske et al. (2017), in which only animals previously habituated to the IA box in the absence of shock reconsolidate the avoidance memory, we simulated the protocol shown in Figure 2A (see Methods). A distinct attractor network represents each experimental group, and, at each stage of memory storage, the groups differ in their synaptic configuration. The matrices in Figure 3A reflect the synaptic configuration after the storage of each memory pattern, including the update by the time-dependent decay factor. In them, the warm colors, closer to yellow, indicate the formation of excitatory synapses, while the cool ones, closer to blue, express inhibitory connections. Since the memories are sparse in relation to the size of the network, a large number of null weights (central green of the color bar) can be seen, indicating the absence of connection between the respective units. After a relatively short reexposure, characterized by a mixed input signal between the shock and non-shock patterns (# = 3.1), the connections formed during training (avoidance memory storage) are mostly reinforced in the vehicle group and weakened in the aniso one. The latter is due to the unique action of the degradation factor since Hebbian learning was blocked by *S* = 0.

The result of the network dynamics was measured at each storage session by retrieval tests from the context input signal (Figure 3B). For each test, a new initial state of the network is randomly generated. In tests performed after the habituation, the median of latencies is slightly higher for the control group due to the related probability density function simulating the lack of knowledge of the IA box. In the same way, the non-shock group simulates habituation in the IA box (i.e., without shock) and the tests after that indicate the retrieval of the non-shock memory pattern, which leads animals to step down faster from the platform since they are already familiar with the environment. On the other hand, after training, which simulates the experimental session in which the animals receive shock after stepping down the platform, the shock memory tends to be retrieved, resulting in long step-down latencies. The networks are then subject to the reexposure stage, which is meant to simulate a short experimental session in which the animal is placed over the platform and subsequently removed from it before stepping down (Radiske et al., 2017). That is, during the reexposure session the animals do not receive a shock, and the network input pattern reflects a conflictual perception of the context (Osan et al., 2011), in which in addition to the shock neurons, the non-shock neurons are also activated, but to a lower extent. Similarly to after training, the shock memory tended to be evoked in retrieval tests performed after reexposure for all vehicle networks and also in aniso networks of the control habituation group. Interestingly, however, only the aniso groups that went through the non-shock habituation (non-shock/aniso) had the behavior altered by the reexposure, in which animals once again exhibited short step-down latencies (Figure 3B). This is because networks of this group mostly retrieved the context-related non-shock habituation memory due to the degradation of the avoidance engram, characterizing the reconsolidation blockade process. Thus, the simulation results obtained here match the experimental findings by Radiske et al. (2017).

The shock attractor was always retrieved in the control habituation group irrespective of aniso since these networks did not form the trace corresponding to the non-shock memory. In the other habituation group, the non-shock engram was previously formed and competed with the shock attractor during retrieval tests since both share the same context neurons. Figure 3C provides an example of the network dynamics for non-shock habituation groups during one retrieval test using the same initial conditions: in the vehicle network, the activity of each unit converges directly to the state corresponding to the avoidance memory pattern, while, for aniso, the network “hesitates” until it converges to the non-shock memory pattern. This can be observed in Figure 3D, where the activity of the shock neurons increases in the first second and then decreases to a latent state, leading to the retrieval of the non-shock pattern.

In order to distinguish the occurrence of the reconsolidation and extinction phenomena, we repeated the previous protocol, but now varying the reexposure input pattern from pure aversive to pure non-aversive (Figure 4A), meant to simulate different durations of the nonreinforced reexposure session (Osan et al., 2011). The result of the post-reexposure retrieval tests is presented in Figure 4B. For the control/vehicle group (blue line in top panel), we found that when the input pattern is close to the non-shock memory (pattern # > 8, representing long reexposure sessions), the networks exhibited extinction, in which the non-shock attractor is successfully stored and subsequently retrieved in post-reexposure tests, leading to short step-down latencies. Noteworthy, we observed that the extinction of the avoidance memory is highly “anticipated” for the non-shock habituation group, in which input patterns associated with shorter sessions already lead to extinction (pattern # > 5, blue line in the bottom panel). For the aniso groups (green lines), the respective input patterns leading to extinction in vehicle networks were instead associated with extinction blockade, in which animals exhibited long step-down latencies in post-reexposure tests. This is because the non-shock memory responsible for the extinction behavior could not be stored in the absence of protein synthesis. For intermediate input patterns preceding the extinction patterns, the aniso groups exhibited reconsolidation blockade as described above (Figure 3B). Interestingly, reconsolidation of the avoidance memory in the non-shock habituation group occurs “before” the same phenomenon in the control habituation group (3 < pattern # < 5 vs. 5 < pattern # < 8). Finally, for input patterns close to the shock memory, representing very short reexposure sessions, neither extinction nor reconsolidation occurred, but only the reinforcement of the pre-existing avoidance memory (in networks with protein synthesis).

We next investigated the influence of Hebbian learning on avoidance memory reconsolidation. To that end, we first assessed, via post-reexposure retrieval tests, the impact of the storage strength of each habituation memory for various reexposure patterns. Figure 5A shows the simulated experimental protocol with the varied parameters highlighted in red, i.e., the *S* factor in habituation and the reexposure input pattern. A sample of results, considering two values of each parameter, is shown in Figure 5B. Note that, for weak habituation storage (*S* = 0.7), there is a reinforcement of the avoidance memory in all habituation groups when the reexposure pattern is closer to the shock one (top left panel), and aniso has no effect. But when the reexposure pattern is closer to the non-aversive engram (top right panel), reconsolidation occurs in the control habituation group along with extinction in the non-shock habituation group, with both phenomena blocked by aniso. However, when the habituation memory is stored more strongly than the training memory (*S* = 1.0), the reexposure pattern closer to an aversive sensation (bottom left panel) results in the reconsolidation of the avoidance memory for the control habituation group and, remarkably, in the extinction of the non-shock memory for the non-shock habituation group. If the reexposure pattern is closer to a non-aversive sensation (bottom right panel), avoidance memory reconsolidation occurs in the control habituation group, and reinforcement of the non-shock memory takes place in the non-shock habituation group.

**Figure 5.**
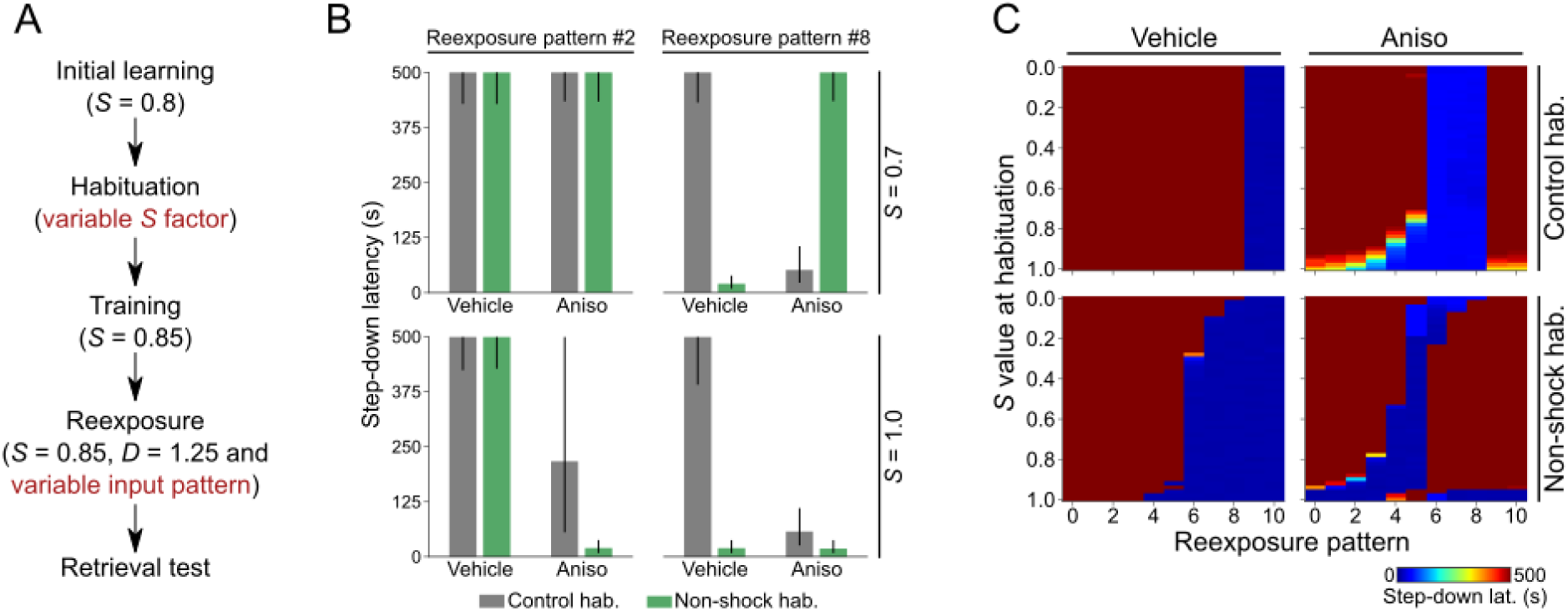
Effect of the habituation memory strength and reexposure pattern in post-reexposure retrieval tests. A) Schematics showing the simulation protocol with the varied parameters in red: habituation *S* factor and reexposure input pattern. B) Step-down latencies in post-reexposure retrieval tests for habituation with *S* value equals to 0.7 and 1.0 and reexposure pattern #2 and #8. For *S* = 0.7 and reexposure pattern #2 (top left), the long step-down latencies indicate the reinforcement of the shock memory during reexposure in all habituation groups. For *S* = 0.7 and reexposure pattern #8 (top right), the step-down latencies indicate the occurrence of shock memory reconsolidation in the control habituation group and extinction in the non-shock habituation group. For *S* = 1.0 and reexposure pattern #2 (bottom left), there is shock memory reconsolidation for control habituation and extinction of the non-shock memory for non-shock habituation. For *S* = 1.0 and reexposure pattern #8 (bottom right), shock memory reconsolidation occurs in the control habituation group and non-shock memory reinforcement in the non-shock habituation group. Results represent the median ± interquartile range of 1000 retrieval tests. C) Color plots summarizing the effect of distinct reexposure patterns and the strength of the habituation memory (*S* value) on post-reexposure retrieval tests. See text for interpretation. Results depict the median of 1000 retrieval tests for each choice of parameters.

The result of the full parameter scan is shown in Figure 5C, where the color plots depict the medians of step-down latencies. In the control/vehicle group (top left panel), the results do not depend on the strength of Hebbian learning during habituation since the control engram is unrelated to the engram stored during training. For this group, the long step-down latencies (red region) from reexposure pattern #0 to #8 indicate reinforcement or reconsolidation of the avoidance memory, while, for patterns above #8, the short latencies (blue region) indicate extinction. When aniso is simulated after reexposure for the control habituation group (top right panel), the results express the blockade of the phenomena observed in vehicle, that is, reconsolidation blockade or extinction blockade (notice that the blockade of reinforcement does not substantially change network behavior). For the non-shock habituation memory, the stronger its storage the more “anticipated” the reexposure pattern is to be enough for reconsolidation or extinction in the vehicle group (bottom left panel) and for their blockade in the aniso group (bottom right panel; see also Figure 4B), with the exception of the networks in which the habituation memory is stored more strongly than the avoidance memory (*S* ≈ 1). In this particular case, extinction of the non-shock memory occurs from pattern #0 to #3, followed by its reconsolidation for patterns #4 and #5 and then reinforcement from pattern #6 to #10.

We then analyzed the impact of the strength of the avoidance memory. Figure 6A shows the schematics of the simulation protocol with the varied parameters in red. The sample results in Figure 6B show that, for a weak training (*S* = 0.7), an aversive reexposure (pattern #2, top left panel) promotes reinforcement of the avoidance memory in the control habituation group. When simulating aniso, such reinforcement does not take place and the step-down latencies do not achieve ceiling values, reflecting a weaker shock attractor than in previous figures. For the non-shock habituation group, we have a similar reinforcement of the shock memory and long step-down latencies for vehicle networks, but for the aniso networks, the non-shock memory attractor dominates and leads to short step-down latencies. This is because the shock memory cannot be properly stored following a single weak training session in this habituation group (not shown), and thus reinforcement during reexposure is required for its attractor to prevail over the previously stored non-shock attractor.

**Figure 6.**
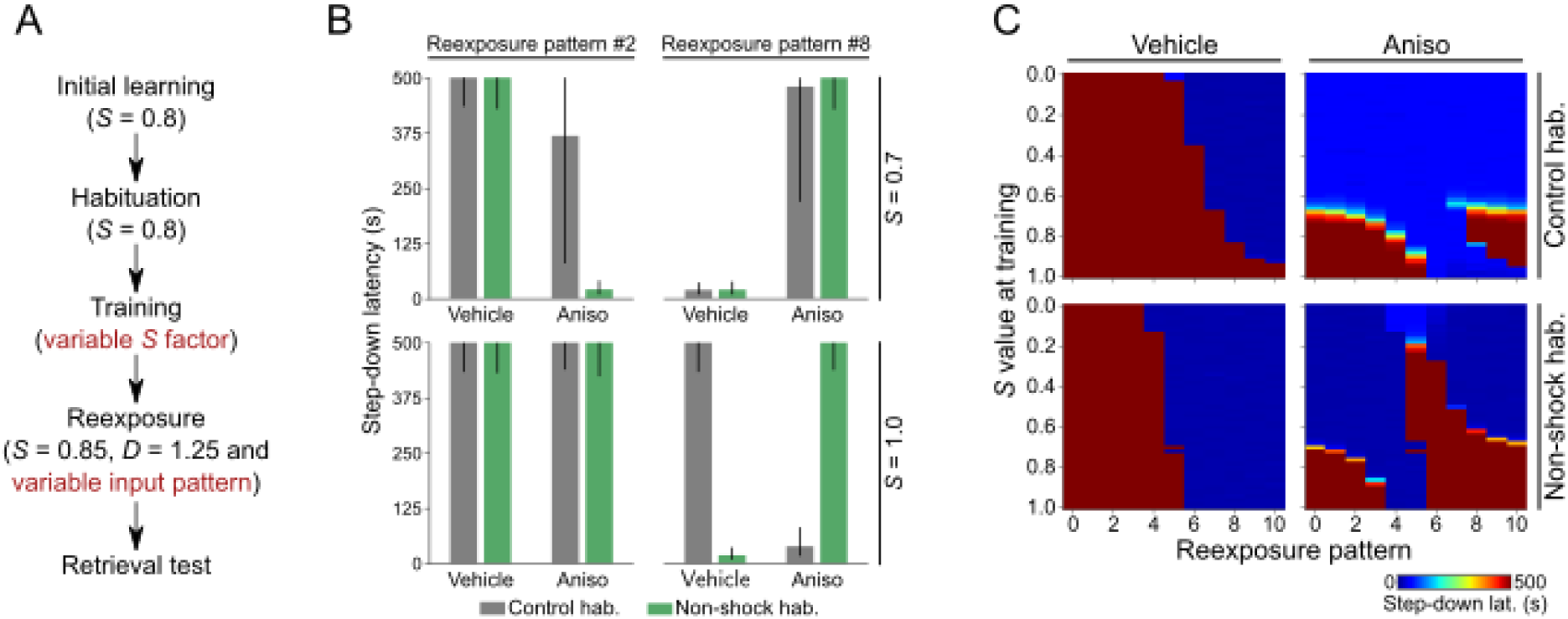
Effect of the training memory strength and reexposure pattern in post-reexposure retrieval tests. A) Schematics showing the simulation protocol with the varied parameters in red: training *S* factor and reexposure input pattern. B) Step-down latencies in post-reexposure retrieval tests for habituation with *S* value equals to 0.7 and 1.0 and reexposure pattern #2 and #8. For S = 0.7, the shock memory is weakly stored, and the reexposure pattern #2 produces its reinforcement for all habituation groups (top left), an effect blocked by aniso (notice that in this case the non-shock memory prevails for the non-shock habituation group). The reexposure pattern #8 (top right) leads to the storage of the non-shock attractor in the control habituation group, and to the reconsolidation of the non-shock attractor in the non-shock habituation group. For S = 1.0 and reexposure pattern #2 (bottom left), there is reinforcement of the shock memory for all habituation groups. For S = 1.0 and reexposure pattern #8 (bottom right), there is shock memory reconsolidation for the control habituation group and its extinction for the non-shock habituation group. Results represent the median ± interquartile range of 1000 retrieval tests. C) Color plots summarizing the effect of distinct reexposure patterns and the strength of the training memory (*S* value) on post-reexposure retrieval tests. See text for interpretation. Results are the median of 1000 retrieval tests for each parameter choice.

On the other hand, reexposure closer to a non-aversive experience (pattern #8, top right panel) leads to the storage of the non-shock attractor in the control habituation group, and, interestingly, to the reconsolidation of the non-shock attractor in the non-shock habituation group. Therefore, notice that the long step-down latencies following aniso in the non-shock habituation group is because of the reconsolidation blockade of the non-shock attractor due to degradation of its synaptic weights, which makes the weakly stored shock attractor now to become dominant. For a strong training (*S* = 1.0), there is reinforcement of the shock memory with the aversive reexposure in all habituation groups (pattern #2, bottom left panel), while the non-aversive reexposure (pattern #8, bottom right panel) leads to shock memory reconsolidation in the control habituation group and extinction in the non-shock habituation group.

Figure 6C shows the full parameter scan. In the vehicle groups (left panels), we found that the stronger the avoidance memory is stored, the more non-aversive the reexposure should be for the occurrence of extinction, with the control habituation group requiring even more non-aversive patterns than the non-shock habituation (cf. the ‘anticipation’ effect mentioned above). Indeed, one reexposure session was insufficient for extinction in the control habituation group when S was close to 1 (top left panel). For the control/aniso group (top right panel) and *S* ≥ 0.7, we observed the blockade of the respective phenomenon taking place in the vehicle group, i.e., blockade of either avoidance memory reinforcement, reconsolidation or extinction for increasing reexposure patterns. For lower *S* values (< 0.7), the aversive memory is stored much weaker than the habituation one, and the latter is always retrieved after an amnesic reexposure. Similar observations hold for the aniso networks in the non-shock habituation group (bottom right panel), that is, for *S* ≥ 0.7 we could sequentially observe reinforcement blockade, reconsolidation blockade or extinction blockade for reexposure patterns approaching the non-aversive pattern. A difference from the control habituation, however, is that for *S* < 0.7 we found a parameter region in which reconsolidation blockade occurred for the non-shock memory instead (compare the top and bottom right panels).

Finally, we analyzed the impact of the degradation factor following reexposure. Figure 7A shows the simulation protocol with the varied parameters in red. The sample results in Figure 7B reveal that, for an aversive reexposure (pattern #2), reinforcement of the avoidance memory occurs for all networks independently of the *D* value (left panels). This is not surprising since synaptic degradation in the model depends on prediction error (see Methods and Osan et al., 2011), while the shock memory and reexposure pattern #2 are quite similar (see Figure 4A); hence, mismatch-induced degradation does not significantly influence network behavior in this case and neither the lack thereof. In the case of a non-aversive reexposure (pattern #8, right panels), the same phenomenon occurs in the control habituation group and low degradation levels (*D* = 0.3; top panel). Nevertheless, for a high degradation level (*D* = 1.4; bottom panel), control habituation networks undergo avoidance memory reconsolidation (notice short step-down latencies indicating post-reexposure amnesia in aniso networks). In the non-shock habituation group, the networks exhibited extinction for both levels of synaptic degradation, which was blocked by aniso.

**Figure 7.**
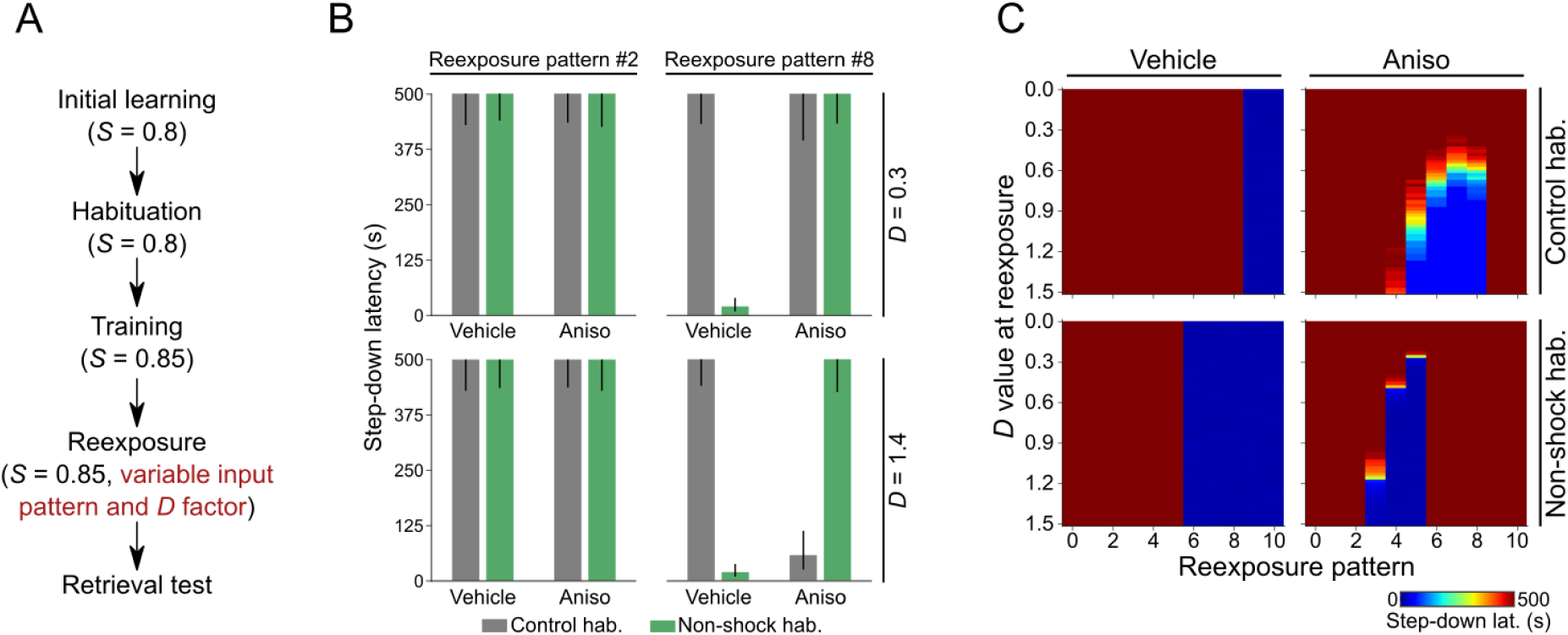
Effect of the synaptic degradation strength and reexposure pattern in post-reexposure retrieval tests. A) Schematics showing the simulation protocol with the varied parameters in red: *D* factor and input pattern in reexposure. B) Step-down latencies in post-reexposure retrieval tests for reexposure with *D* value equal to 0.3 and 1.4 and reexposure pattern #2 and #8. For the reexposure pattern #2 (left), *D* does not affect the behavioral outcome, and shock memory reinforcement occurs. For the reexposure pattern #8 with *D* = 0.3 (top right), there is shock memory reinforcement for the control habituation group and extinction for the non-shock habituation. When *D* = 1.4 (bottom right), there is reconsolidation of the aversive memory in the control habituation group and its extinction in the non-shock habituation group. Results represent the median ± interquartile range of 1000 retrieval tests. C) Color plots summarizing the effect of distinct reexposure patterns and the strength of the synaptic degradation (*D* value) on post-reexposure retrieval tests. For the vehicle networks (left panels), the synaptic degradation level does not influence behavioral outcomes. For aniso networks, the higher the degradation level, the larger the window for avoidance memory reconsolidation to occur. Notice further that aversive memory extinction (vehicle) and extinction blockade (aniso) are not affected by changes in *D*. Results depict the median of 1000 retrieval tests for each parameter choice.

As shown in Figure 7C, the full parameter scan reveals that synaptic degradation directly influences avoidance memory reconsolidation, where the chance of its occurrence is proportional to *D*. In other words, the higher the level of synaptic degradation, the larger the reconsolidation “window” (i.e., the number of reexposure patterns in which reconsolidation occurs), as can be observed in the aniso networks (right panels). Moreover, as in the previous analyses, the reconsolidation window is anticipated for the non-shock habituation. Interestingly, extinction (vehicle networks) and extinction blockade (aniso networks) differ between habituation groups (see Figure 4) but do not depend on the synaptic degradation level. Mechanistically, this is because extinction depends only on Hebbian learning of the non-shock attractor, which is influenced by the factor *S* but not *D*.

## Discussion

The attractor network model of Osan et al. (2011) simulates the processes of pattern completion and separation observed, for example, during the retrieval of hippocampal-dependent memories (Guzowski et al., 2004; Neunuebel and Knierim, 2014). Moreover, its properties provide a mapping of computational variables into biological substrates in a particularly simple and acceptable way, which makes it an attractive tool to study the phenomena of memory reconsolidation and extinction. By successfully simulating the effect of boundary conditions, the model by Osan et al. (2011) supports the hypothesis that reconsolidation and extinction use the same reinforcement and synaptic stabilization system, being only distinct instances of the same process (Almeida-Corrêa and Amaral, 2014). Unlike other models, the parametric distinction between Hebbian learning and synaptic degradation induced by prediction error enables analyses that mimic the action of pharmacological agents that differentially affect them (Lee et al., 2008; Gershman et al., 2017).

Although the model does not have a hierarchical structure of message exchange, it can be considered a way of predictive coding (Friston, 2018; Rao and Ballard, 1999) due to its recurrent processing structure (Pennartz et al., 2019). Here, learning and perception participate in the same process of modeling the world, characterizing the brain as an inference machine by predicting and explaining its sensations and also adapting itself to minimize prediction errors by adjusting its synaptic structure, which leads the model to be also in accordance with the free-energy principle (Friston, 2010; Friston et al., 2006). The model thus tries to explain memory reconsolidation and extinction in a simple structural perspective. It should be noted, however, that the model does not address the theory that there is a null point between the phenomena (Cassini et al., 2017; Merlo et al., 2014), i.e., the transition between reconsolidation and extinction is abrupt in terms of network structure and its dynamics.

Simulating reexposure to a previously learned stimulus/context associated with an aversive sensation leads the network to different possible states depending on the degree of similarity between the original experience and the new one. A stimulus identical to that already learned causes the reinforcement of the aversive memory engram, and such effect is blocked when one of the molecular requirements of synaptic plasticity, such as protein synthesis, is inhibited. Of note, considering that memory formation results from a balance among the synthesis and degradation of synaptic proteins (Park and Kaang, 2019), in the phenomenon we call reinforcement here, the strengthening of connections stands out in relation to synaptic degradation.

When the input signal moderately differs from the original trace, the latter is restored from a new configuration of synaptic weights from the balance between Hebbian learning and synaptic degradation induced by prediction error. In this case, when Hebbian plasticity is blocked, the effect of synaptic degradation prevails, causing the weakening of the previously stored avoidance memory and, consequently, reconsolidation blockade. However, if the input signal associated with the reexposure session substantially differs from the original engram, the formation of a new memory trace takes place (extinction), establishing also inhibitory associations between the context and aversive engrams, leaving the latter in a latent state. Such extinction based on Hebbian learning of the non-aversive engram can also be blocked by inhibition of protein synthesis, preserving the initial aversive memory pattern (Almeida-Corrêa and Amaral, 2014). Of note, in Figure 4B, the reexposure “windows” called reconsolidation and extinction are study targets for the treatment of fear-related disorders.

Here we investigate whether the formalism introduced by Osan et al. (2011) is general enough to explain the behavioral results of Radiske et al. (2017) in the inhibitory avoidance (IA) paradigm, in which the previous learning of the IA box as non-aversive was shown to be a boundary condition for the reconsolidation of the avoidance memory. Consistent with their results, our simulations show that for certain patterns of cortical afference during nonreinforced reexposure, the prior knowledge of the context as non-aversive causes amnesic behavior in networks subjected to inhibition of Hebbian learning. In these cases, the storage of the non-shock memory attractor during habituation leads to its later retrieval in post-reexposure tests following mismatch-induced degradation of the shock memory trace upon conflictual reexposures. The latter is caused by the divergence between the neural network input signal (a mixture of shock and non-shock patterns) and the recalled engram upon reexposure (the shock pattern), which characterizes the prediction error. Even when the habituation memory is unrelated to the training context, such divergence may still occur, weakening the shock memory trace. But for networks habituated with the control (“open field”) memory, the reconsolidation of the avoidance memory requires a greater divergence between the input signal and the shock memory (see Figure 4), which, in turn, is associated with longer nonreinforced reexposures to the context (IA box). However, as opposed to contextual fear conditioning, relating the time of nonreinforced reexposure in the IA paradigm with the animal’s representation/perception of the context is difficult because, in the work of Radiske et al. (2017), the rats do not step down from the platform during the 40 seconds of reexposure, so they do not experience the lack of shock on the ground.

Radiske et al. (2017) interpreted that the behavioral distinction between the different habituation groups was due to the non-participation of the hippocampus in the reexposure of the “control” habituation group since the prediction error would not occur there. In other words, the hippocampus would participate in the reconsolidation of the avoidance memory only when animals were subjected to the non-shock habituation, i.e., when pre-exposed to the training environment without shock before acquiring the aversive trace related to such context — an idea that is corroborated by the analysis of hippocampal theta and gamma oscillations (Radiske et al., 2017). From a computational standpoint, for a reexposure input pattern in which there is little prediction error, the attractor network proposed by Osan et al. (2011) acts to reinforce the avoidance memory trace in the control habituation group and to reconsolidate it in the non-shock habituation group. Thus, the model by Osan et al. (2011) suggests that the result of Radiske et al. (2017) can be explained without assuming that different structures are activated due to prior habituation.

The results of the parameter scan provide experimental predictions for the IA paradigm. Specifically, the model predicts that if rats were reexposed to the context for a longer duration without the shock (so that they could step down the platform), for a same reexposure duration animals in the control habituation group could exhibit reconsolidation of the aversive memory, while those of non-shock habituation could display extinction (e.g., Figures 4 and 5). Therefore, the results further suggest that the approach to be taken in exposure therapy for the treatment of phobias and fear-related disorders (McNally, 2007) depends on whether or not the subject has prior knowledge of the context, currently aversive, as non-aversive. Thus, blocking reconsolidation during nonreinforced reexposures with amnesic drugs could be a treatment better applied to the group that, in the first contact with a new environment or context, relates it to the aversive sensation experienced there. On the other hand, when such an environment or context has been previously experienced as non-aversive, exposure therapy without medication could be better utilized to induce extinction. In the first case, it seems to be easier to block the reinforcement of the connections between the context neurons and the aversive engram, letting the synaptic degradation act towards weakening the trace. In the second, the safe memory trace can more easily prevail with long nonreinforced reexposures, leading to the extinction of the aversive memory. Generally, the simulations reveal that the previous knowledge of a safe context influences the width of the occurrence windows of each phenomenon. While for the control habituation group there are large reexposure windows for reinforcement and reconsolidation of the avoidance memory, these windows are reduced in the non-shock habituation group, with more opportunity for extinction incidence (see Figures 4 to 7).

To test the model predictions, we propose the conduction of new experiments with animals exposed to different reexposure times. Thus, the animals would be subjected to various levels of perception (reexposure input pattern), which, according to the model, would influence their behavior during retrieval testing. In parallel, for such conditions, we suggest empirically investigating the influence of the strength of the habituation and training memories (e.g., by changing exploration times or shock amplitude), as well as the influence of enhancers and inhibitors protein synthesis and synaptic degradation to better map their behavioral effects.

Finally, although this paper deals with the case of reconsolidation and extinction of avoidance memory in the IA paradigm, it more generally highlights the balance between Hebbian learning and synaptic degradation as a key factor in modulating such phenomena. These theoretical insights can also be applied in other scenarios involving the learning of aversive memories followed by nonreinforced reactivations. In this sense, the simulation of the behavior (here, step-down latency) in retrieval tests is rather a detail behind the big picture of the network dynamics supported by the model.

## Acknowledgments

This study was supported by CNPq and CAPES (Finance Code 001), Brazil. We thank A. Radiske, M.C. Gonzalez, J.I. Rossato, O. B. Amaral, and R. N. Leão for critical discussions.

